# Distinct sensitivities to SARS-CoV-2 variants in vaccinated humans and mice

**DOI:** 10.1101/2022.02.07.479468

**Authors:** Alexandra C. Walls, Laura A. VanBlargan, Kai Wu, Angela Choi, Mary Jane Navarro, Diana Lee, Laura Avena, Daniela Montes Berrueta, Minh N. Pham, Sayda Elbashir, Marcos C. Miranda, Elizabeth Kepl, Max Johnson, Alyssa Blackstone, Kaitlin Sprouse, Brooke Fiala, Megan A. O’Connor, Natalie Brunette, Prabhu S. Arunachalam, Lisa Shirreff, Kenneth Rogers, Lauren Carter, Deborah H. Fuller, Francois Villinger, Bali Pulendran, Michael S. Diamond, Darin K. Edwards, Neil P. King, David Veesler

## Abstract

The emergence of severe acute respiratory syndrome coronavirus 2 (SARS-CoV-2) in 2019 has led to the development of a large number of vaccines, several of which are now approved for use in humans. Understanding vaccine-elicited antibody responses against emerging SARS-CoV-2 variants of concern (VOC) in real time is key to inform public health policies. Serum neutralizing antibody titers are the current best correlate of protection from SARS-CoV-2 challenge in non-human primates and a key metric to understand immune evasion of VOC. We report that vaccinated BALB/c mice do not recapitulate faithfully the breadth and potency of neutralizing antibody responses against VOC, as compared to non-human primates or humans, suggesting caution should be exercised when interpreting data for this animal model.

---

Vaccine candidates are typically evaluated in small mammals (e.g. mice) and non-human primates (NHPs) prior to cGMP manufacturing and human clinical trials. For SARS-CoV-2, serum neutralizing antibody titers represent the current best correlate of protection in NHP challenge studies^1,2^ and in humans^3^. Serum neutralizing antibody titers are also used as metrics in human clinical trials to benchmark new vaccine candidates (e.g. NCT05007951 comparing GBP510 to AZD1222) and the administration of neutralizing monoclonal antibodies has been shown to improve disease outcome for some COVID-19 patients^4^. As variants of SARS-CoV-2 emerge^5–9^ it is necessary to evaluate their impact on serum neutralizing activity, as a proxy for vaccine efficacy, in order to inform public health policies worldwide and further vaccine development.

We compared neutralizing antibody responses elicited following vaccination of BALB/c mice with three distinct AddaVax-adjuvanted protein subunit immunogens. Mice were immunized at week 0 and 3 with a clinical-stage, multivalent RBD-nanoparticle (RBD-NP)^1,10,11^, an S ‘2P’ prefusion-stabilized S^10,12^, or a next-generation prefusion-stabilized HexaPro S^11,13^. Sera were collected 2, 5, and 8 weeks post-prime and serum neutralizing activity (expressed as the dilution inhibiting 50% of entry: ID_50_) was evaluated using single-round vesicular stomatitis virus (VSV) pseudotyped with G614 S, Beta S (B.1.351: L18F, D80A, D215G, 17mccaL242-L244 deletion, R246I, K417N, E484K, N501Y, D614G, A701V), or Gamma S (P.1: L18F, T20N, P26S, D138Y, R190S, K417T, E484K, N501Y, D614G, H655Y, T1027I) in two distinct cell lines (VeroE6/TMPRSS2^14^ and HEK293T/ACE2^15^).

Of the three immunogens tested, only RBD-NP induced detectable serum neutralizing activity post-prime (2 weeks post immunization), in contrast to what we observed with AS03-adjuvanted HexaPro S in NHPs^1,10^. With RBD-NP-elicited sera at week 2, we observed a 2.0-fold reduction and an equivalent neutralization potency against the Beta and Gamma variants of concern (VOC) relative to G614 using VeroE6-TMPRSS2 cells, respectively **(Figure 1A, Figure S1)** and 2.9- and 1.3-fold decreases using HEK293T-ACE2 cells, respectively **(Figure 1B, Figure S2)**. All three protein immunogens induced robust neutralizing antibody responses 2 weeks post-boost. Serum neutralizing activity relative to G614 at week 5 was increased 2.4-fold (Beta) and decreased 1.2-fold (Gamma) in mice immunized with RBD-NP, reduced 1.5-fold (Beta) and increased 2.0-fold (Gamma) for S ‘2P’, and reduced 1.2-fold (Beta) and increased 1.4-fold (Gamma) with HexaPro S-elicited sera using VeroE6-TMPRSS2 cells **(Figure 1A, Figure S1)**. These findings were overall consistent with the outcomes of the assays using HEK293T-ACE2 cells **(Figure 1B, Figure S2)**, with the exception of neutralizing antibody responses elicited by HexaPro S, which were reduced 5-fold against Beta and 2.8-fold against Gamma relative to G614, indicating that these are not antigen-or cell type-specific observations.

**Figure 1:**
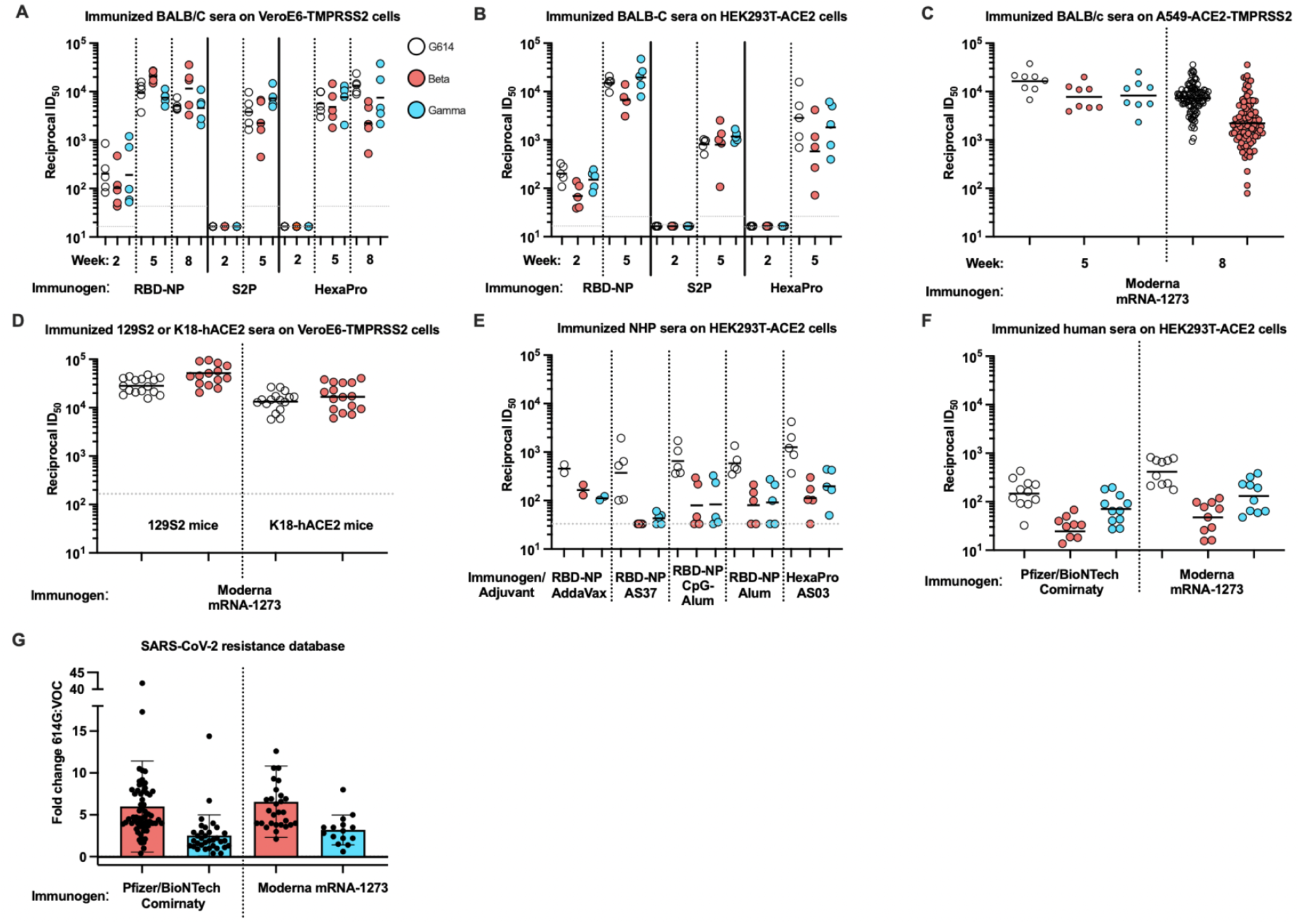
Vaccine-elicited serum neutralizing activity against VOC differs in mice relative to NHPs and humans. (**A**) Neutralization of sera from BALB/c mice immunized with RBD-NP or HexaPro 2 weeks post-prime, 2 weeks post-boost (5 weeks post-prime), or 5 weeks post-boost (8 weeks post-prime) with VSV pseudotyped with SARS-CoV-2 G614 S, Beta S, or Gamma S on VeroE6-TMPRSS2 cells. Data from one out of at least two representative experiments shown; n=5 mice. (**B**) Neutralization of sera from BALB/c mice immunized with RBD-NP or HexaPro 2 weeks post-prime or 2 weeks post-boost (5 weeks post-prime) with VSV pseudotyped with SARS-CoV-2 G614 S, Beta S, or Gamma S on HEK-293T-ACE2 cells. Data from one out of at least two representative experiments shown; n=5 mice. (**C**) Neutralization of sera from BALB/c mice immunized with mRNA-1273 2 or 5 weeks post boost with VSV pseudotyped with SARS-CoV-2 G614 S, Beta S, or Gamma S on A549-ACE2-TMPRSS2 cells. Data from one out of at least two representative experiments shown; n=8 or 104 mice. (**D**) Neutralization of sera from 129S2 or K18-hACE2 mice immunized with mRNA-1273 3 weeks post-boost with authentic SARS-CoV-2 G614 or Beta on VeroE6-TMPRSS2 cells. Data shown are from two independent experiments; n=16 mice. (**E**) Neutralization of sera from NHPs immunized with RBD-NP formulated in various adjuvants or HexaPro at peak titer (day 42 post-prime) with VSV pseudotyped with SARS-CoV-2 G614 S, Beta S, or Gamma S on HEK-293T-ACE2 cells. Data from one out of at least two representative experiments shown; n=2 pigtail macaques, n=5 rhesus macaques. (**F**) Neutralization of sera from humans vaccinated either with Pfizer/BioNTech Comirnaty or Moderna mRNA-1273 on HEK-293T-ACE2 cells (**Table S1**). Data from one out of at least two representative experiments shown; n=10 Moderna mRNA-1273 vaccines, n=11 Pfizer/BioNTech Comirnaty vaccines. Normalized curves and data fits for **A-F** are shown in **Figure S1-6**.(**G**) Fold change data from the SARS-CoV-2 resistance database (https://covdb.stanford.edu/page/susceptibility-data/) for human sera vaccinated with either Pfizer/BioNTech Comirnaty or Moderna mRNA-1273 and assayed with G614 compared to either Beta (n=62 Pfizer/BioNTech Comirnaty, n=29 Moderna mRNA1273) or Gamma (n=34 Pfizer/BioNTech Comirnaty, n=15 Moderna mRNA-1273).

To further understand the generalizability of these observations, we immunized BALB/c mice twice (3 weeks apart) with mRNA-1273^16^ and collected sera at week 5. Serum neutralizing activity 2 weeks post-boost, measured in a different laboratory, was reduced 2.3-fold against both Beta and Gamma variant pseudoviruses compared to G614 S using A549/ACE2/TMPRSS2 cells **(Figure 1C, Figure S3)**^16^. To determine whether these findings were specific to BALB/c mice, we immunized K18-hACE2 transgenic and 129S2 mice with mRNA-1273 twice (3 weeks apart) and collected sera at week 6^17^. Serum neutralizing activity was evaluated using a focus-reduction neutralization test (FRNT) with authentic SARS-CoV-2 carrying the G614 S glycoprotein in a WA1/2020 background or a clinical isolate of the Beta VOC^18^. We observed a 1.8-fold and a 1.25-fold increased neutralization potency of the Beta variant compared to G614 for 129S2 and K18-hACE2 immunized mice, respectively (**Figure 1D, Figure S4**). These data indicate that the unexpected apparent resilience of variants to polyclonal neutralizing antibody responses in mice is independent of the mouse model, vaccine platform, the nature of the antigen, the neutralization assay used or the target cell type.

To understand whether these observations extended beyond mice, we analyzed sera obtained from pigtail macaques and rhesus macaques immunized with RBD-NP or HexaPro S protein subunit vaccines^1,10,11^. Pigtail macaques (n=2) were immunized with AddaVax-adjuvanted RBD-NP at weeks 0 and 4, and sera from week 12 were analyzed. Rhesus macaques were immunized at weeks 0 and 3 with RBD-NP adjuvanted with AS37 (n=5), CpG-Alum (n=5), or Alum (n=5), or with HexaPro S adjuvanted with AS03 (n=6), and sera from week 6 were used. All NHP serum neutralizing titers were lower than immunogen-matched mice titers. Serum neutralizing activity from pigtail macaques immunized with AddaVax-adjuvanted RBD-NP showed a 2.7-fold potency dampening against Beta and 4-fold against Gamma VOC compared to G614 S pseudovirus using HEK293T-ACE2 cells **(Figure 1E, Figure S5)**. Rhesus macaques immunized with AS37-adjuvanted RBD-NP had a reduction in neutralization potency of ≥ 11-fold (Beta) and 8.6-fold (Gamma), whereas a similar ∼6-8-fold decrease was detected with CpG-Alum and Alum for both Beta (8.2 for CpG-Alum and 7.2 for Alum) and Gamma (7.8 for CpG-Alum and 6.4 for Alum) using HEK293T-ACE2 cells **(Figure 1E, Figure S5)**. AS03-adjuvanted HexaPro S immunization led to 11.2- and 6.4-fold reductions in serum neutralizing activity against Beta and Gamma, respectively **(Figure 1E, Figure S5)**. These data recapitulate previous findings made with authentic Beta virus neutralization assays^1^, suggesting that these outcomes are not specific to pseudovirus assays. Collectively, these data obtained with different clinical adjuvants indicate that vaccine-elicited neutralizing antibody responses in NHPs are significantly affected by VOCs, unlike those in BALB/c mice.

Analysis of serum neutralizing antibody titers in individuals vaccinated with either Pfizer/BioNTech Comirnaty or Moderna mRNA-1273 confirmed that the magnitudes of potency reduction against the Beta and Gamma variants relative to G614 is on par with our NHP data and that both are much greater than those observed in BALB/c mice **(Figure 1F, Figure S6, Table S1)**. Compared to G614 S, we observed 6.1-(Beta) and 2.1-(Gamma) fold reductions in neutralization potency with a cohort of humans immunized with Pfizer/BioNTech Comirnaty and 8.7-(Beta) and 3.2-(Gamma) fold reductions with a cohort of humans immunized with Moderna mRNA-1273 using a VSV pseudovirus assay with HEK293T/ACE2 target cells (**Figure 1F, Figure S6)**. These results are in line with the average attenuations of neutralizing antibody responses retrieved from the coronavirus antiviral and resistance database (https://covdb.stanford.edu/), which are 7.3 (Beta) and 4.1 (Gamma) for Pfizer/BioNTech Comirnaty and 7.8 (Beta) and 4.5 (Gamma) for Moderna mRNA-1273 **(Figure 1G)**.

Vaccine-elicited BALB/c mice sera post-prime and post-boost did not recapitulate observations made with NHPs and humans regarding the impact of SARS-CoV-2 variants on neutralizing activity regardless of overall titer, vaccine modality, cell type, or antigen. We next tested whether using sera from mice 8 weeks post-prime would reveal larger differences between G614 and VOC. At 8 weeks post-RBD-NP prime (5 weeks post boost), mouse serum neutralizing activity remained mildly affected when comparing G614 to either Beta or Gamma S VSV pseudotyped viruses in VeroE6/TMPRSS2 cells (**Figure 1A, Figure S1**). A mild attenuation (3.3-fold) of serum neutralizing activity was also observed 8 weeks post-prime against the Beta variant with a cohort of 104 BALB/c mice immunized with mRNA-1273 (**Figure 1C, Figure S3**). In contrast, HexaPro S-immunized mice exhibited a 6-fold drop of serum neutralizing activity against Beta and 1.3-fold against Gamma S VSV pseudotyped viruses (**Figure 1A, Figure S1**), more closely resembling the expected trends for NHP and human data (**Figure 1E-F, Figure S5-6**).

Although BALB/c and other mice are widely used for evaluating vaccine candidates, our data suggest that they are not always an adequate animal model to evaluate the breadth and potency of neutralizing antibody responses against emerging SARS-CoV-2 VOC, potentially due to their different immune repertoires compared to primates. For example, polyclonal neutralizing responses resulting from infection or vaccination with ancestral SARS-CoV-2 isolates in humans and NHPs are hyperfocused on position 484 in the RBD, which does not appear to be recapitulated in mice^11,19^. However, intranasal delivery of an S ‘2P’-based chimpanzee adenovirus-vectored vaccine (ChAd-SARS-CoV-2-S) in K18-hACE2 transgenic mice^20^ and intramuscular delivery of SARS-CoV-2 S mRNA-LNP to BALB/c mice^21^ both show responses in line with the expected human response from G614 and the Beta VOC, suggesting there could be exceptions to the generalizability of these findings. Further experiments will be necessary to understand the immunological basis of these findings, such as comparing vaccine-elicited responses in inbred vs. outbred mouse strains^22^ or the microbial environment of the mice^23^, as our data suggest that other animal models may be better suited to evaluate immune evasion in new variants as they emerge, even though mice are a cost- and time-effective option for evaluating overall immunogenicity.

## Author contributions

A.C.W., and D.V. conceived the project and designed experiments. A.C.W., L.A.V., K.W., A.C., M.J.N., D.L., L.A., D.M.B., S.E., and K.S. performed experiments. M.N.P., M.C.M., E.K., M.J., A.B., B.F., M.A.O., N.B., P.S.A., L.S., K.R., L.C., D.H.F., F.V., and B.P. prepared immunogens and coordinated immunizations. A.C.W., D.H.F. M.S.D., D.K.E., N.P.K., D.V. supervised the project and obtained funding. A.C.W. and D.V. analyzed the data and wrote the manuscript with input from all authors.

## Acknowledgements

We thank Hideki Tani (University of Toyama) for providing the reagents necessary for preparing VSV pseudotyped viruses and Wesley Van Voorhis for coordination of human sera samples. This study was supported by the National Institute of Allergy and Infectious Diseases (DP1AI158186 and HHSN272201700059C to D.V., R01 AI157155 to M.S.D., and Influenza Research and Response (CEIRR) contract 75N93021C00014), a Pew Biomedical Scholars Award (D.V.), an Investigators in the Pathogenesis of Infectious Disease Awards from the Burroughs Wellcome Fund (D.V.), Fast Grants (D.V.), and the Bill & Melinda Gates Foundation (OPP1156262 to N.P.K. and D.V.). D.V. is an Investigator of the Howard Hughes Medical Institute.

## Competing Interests

A.C.W., N.P.K., and D.V. are named as inventors on patent applications filed by the University of Washington based on the RBD-NP presented in this paper. N.P.K. is a co-founder, shareholder, paid consultant, and chair of the scientific advisory board of Icosavax, Inc. and the King lab has received an unrelated sponsored research agreement from Pfizer. D.V. has received an unrelated sponsored research agreement from Vir Biotechnology, Inc. M.S.D. is a consultant for Inbios, Vir Biotechnology, and Carnival Corporation, and on the Scientific Advisory Boards of Moderna and Immunome. The Diamond laboratory has received unrelated funding support in sponsored research agreements from Vir Biotechnology, Kaleido, and Emergent BioSolutions and past support from Moderna not related to these studies. K.W., A.C., and D.K.E are employees of Moderna and hold stock/stock options in the company. D.H.F. has equity interest in HDT Bio.

**Table S1:** Demographics of human sera samples

| Study ID | Age | Vaccine Type | Days post second vaccine | M/F |
| --- | --- | --- | --- | --- |
| 136024 | 65 | Pfizer | 10 | M |
| 136025 | 55 | Pfizer | 18 | M |
| 136026 | 42 | Pfizer | 9 | F |
| 136028 | 63 | Pfizer | 10 | M |
| 136029 | 27 | Pfizer | 8 | F |
| 136031 | 37 | Pfizer | 21 | F |
| 136033 | 62 | Pfizer | 15 | M |
| 136034 | 54 | Pfizer | 14 | F |
| 136036 | 32 | Pfizer | 13 | F |
| 136037 | 52 | Pfizer | 11 | M |
| 136039 | 32 | Pfizer | 22 | F |
| 136032 | 36 | Moderna | 7 | M |
| 136040 | 40 | Moderna | 20 | M |
| 136041 | 64 | Moderna | 16 | M |
| 136042 | 34 | Moderna | 23 | F |
| 136045 | 35 | Moderna | 20 | M |
| 136046 | 40 | Moderna | 24 | M |
| 136047 | 55 | Moderna | 20 | M |
| 136050 | 36 | Moderna | 27 | F |
| 136051 | 53 | Moderna | 20 | F |
| 136052 | 47 | Moderna | 21 | M |

**Figure S1:**
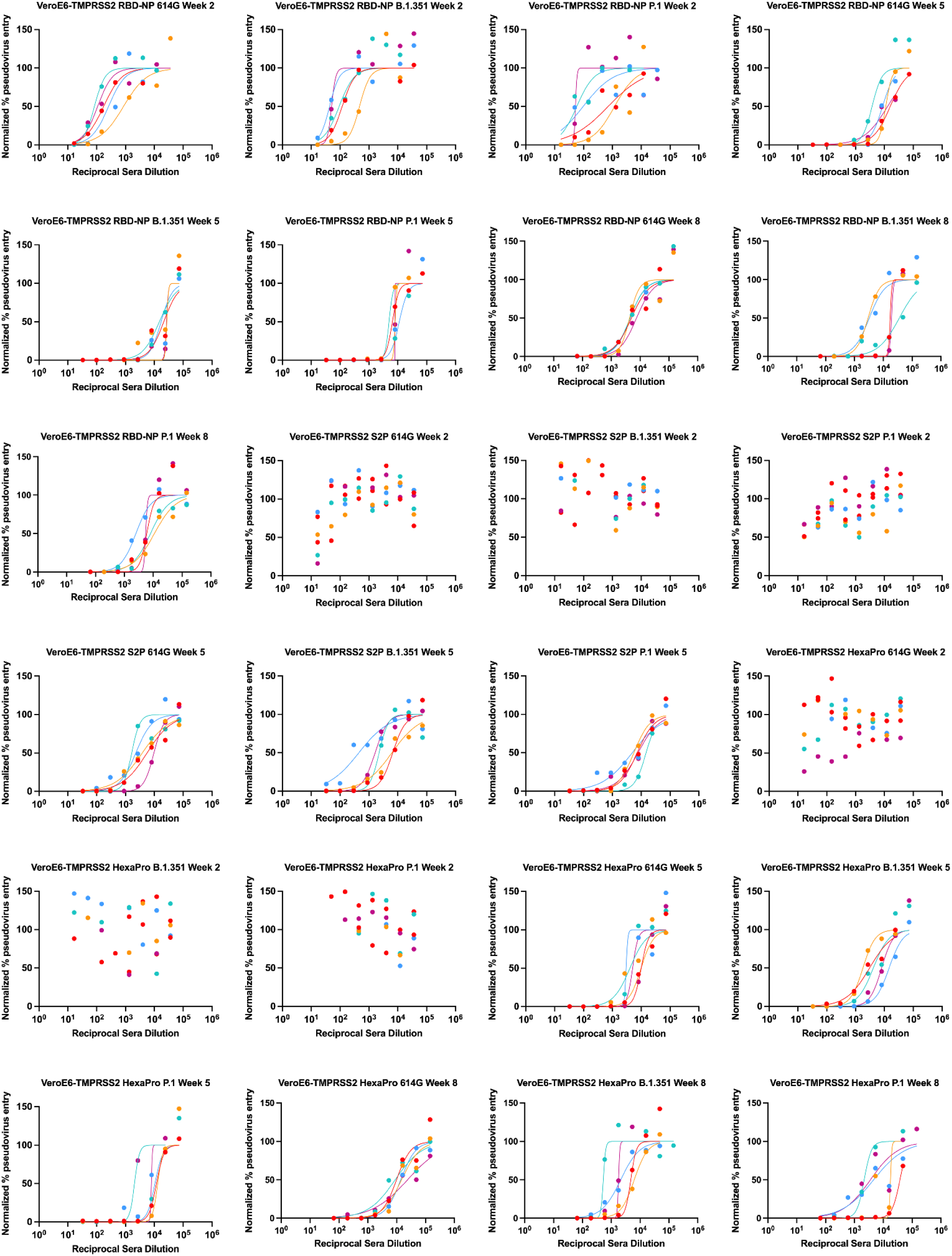
Normalized neutralization curves relating to Figure 1A.

**Figure S2:**
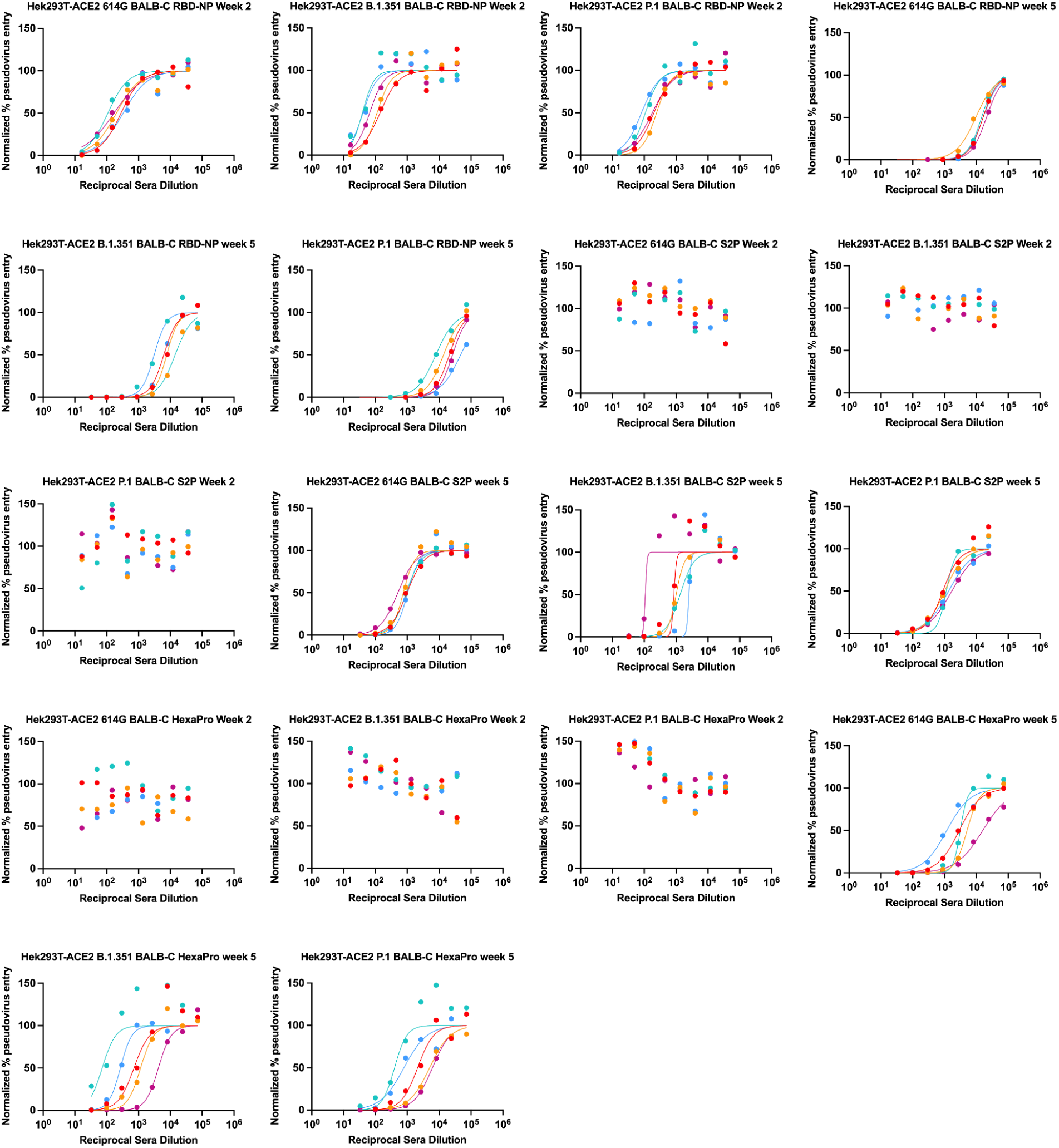
Normalized neutralization curves relating to Figure 1B.

**Figure S3:**
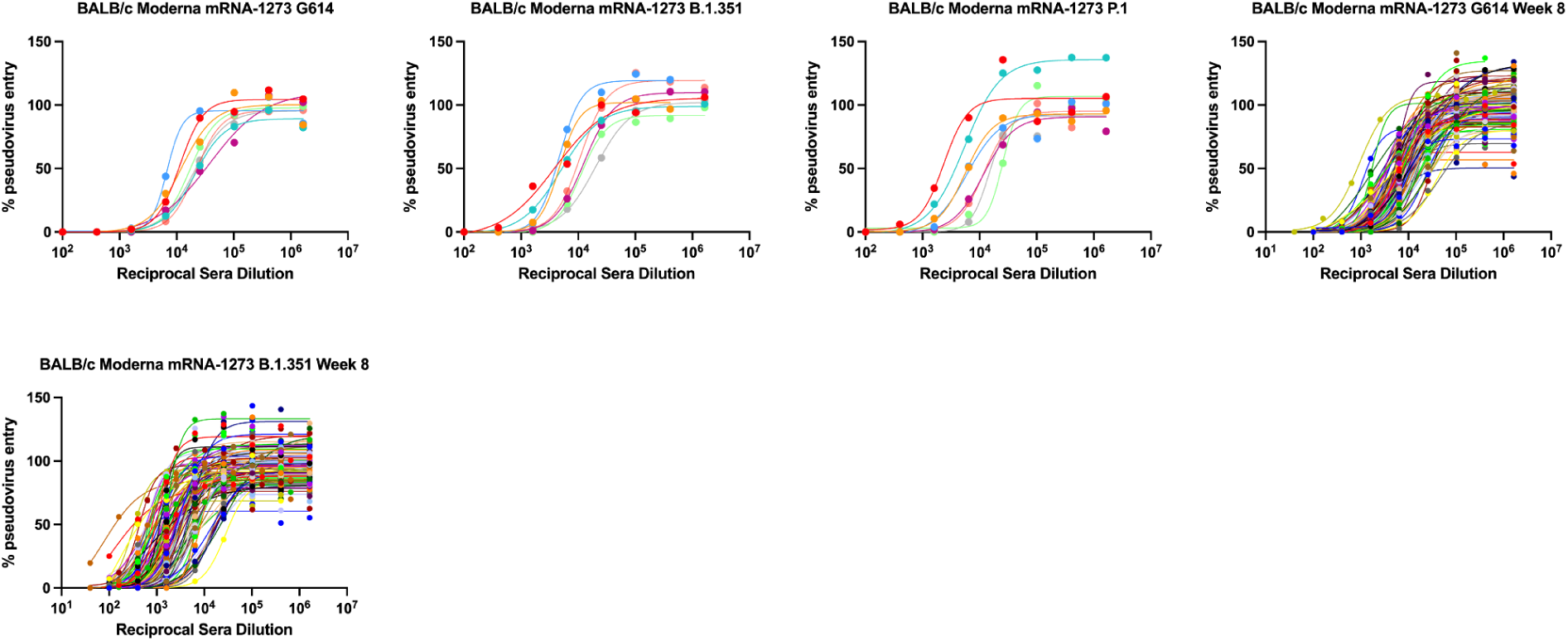
Normalized neutralization curves relating to Figure 1C.

**Figure S4:**
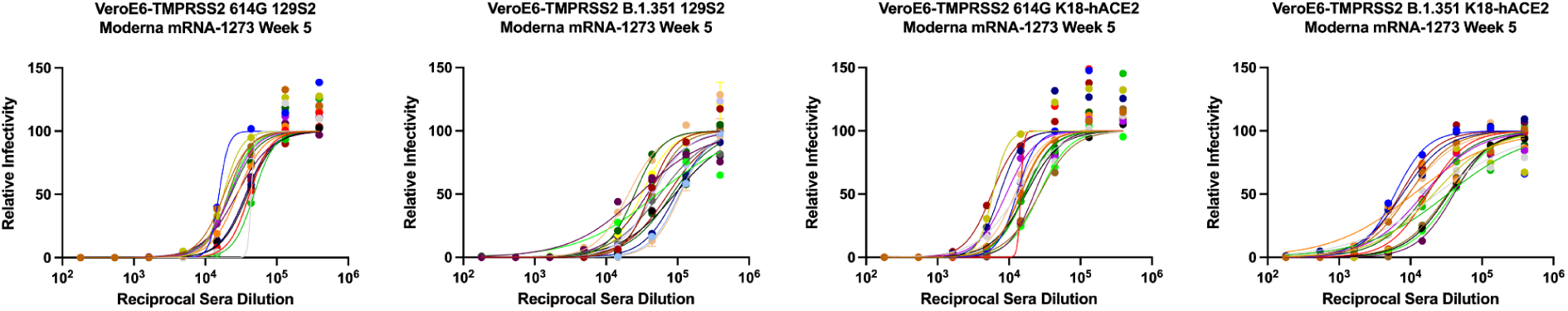
Normalized neutralization curves relating to Figure 1D.

**Figure S5:**
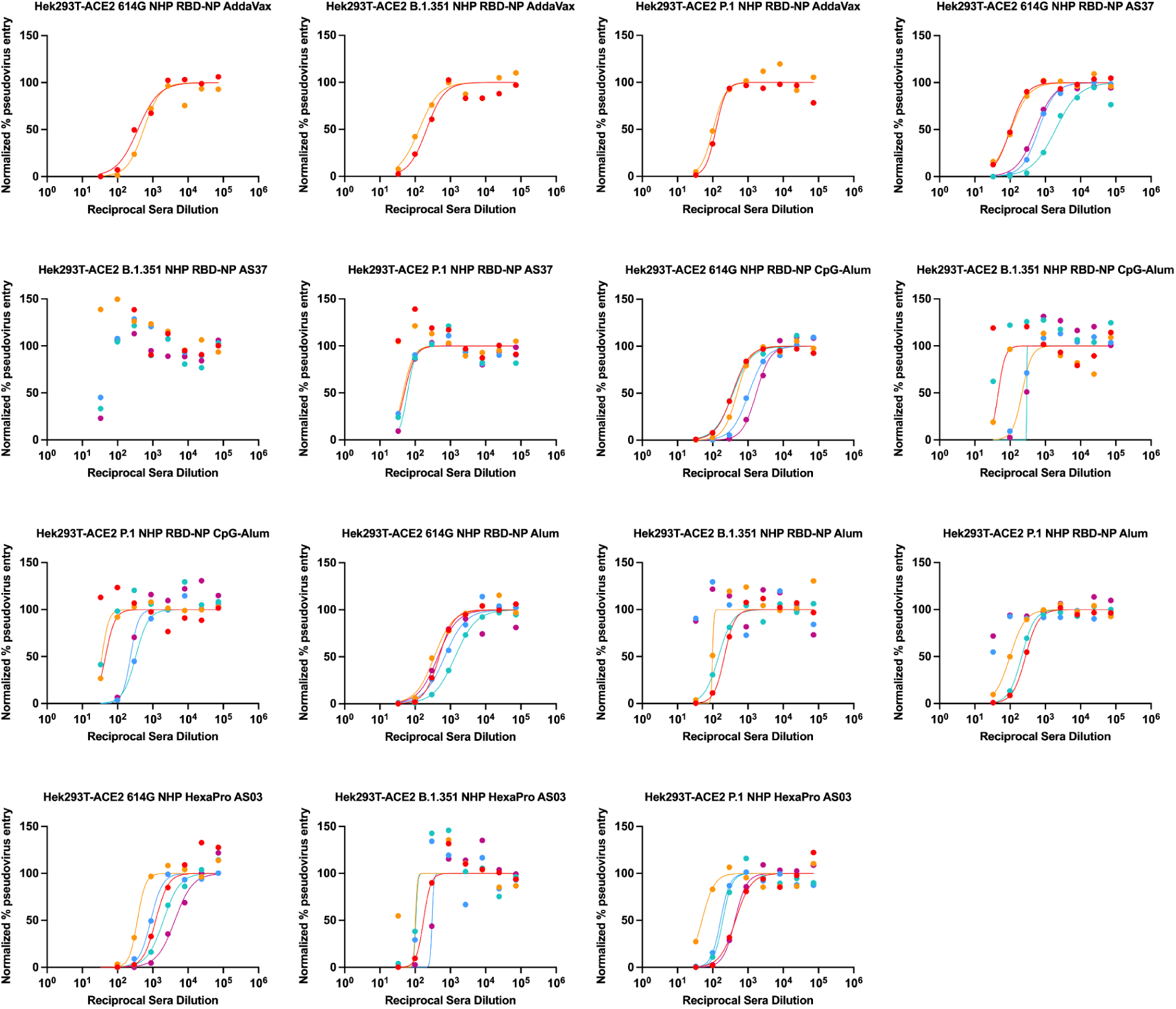
Normalized neutralization curves relating to Figure 1E.

**Figure S6:**
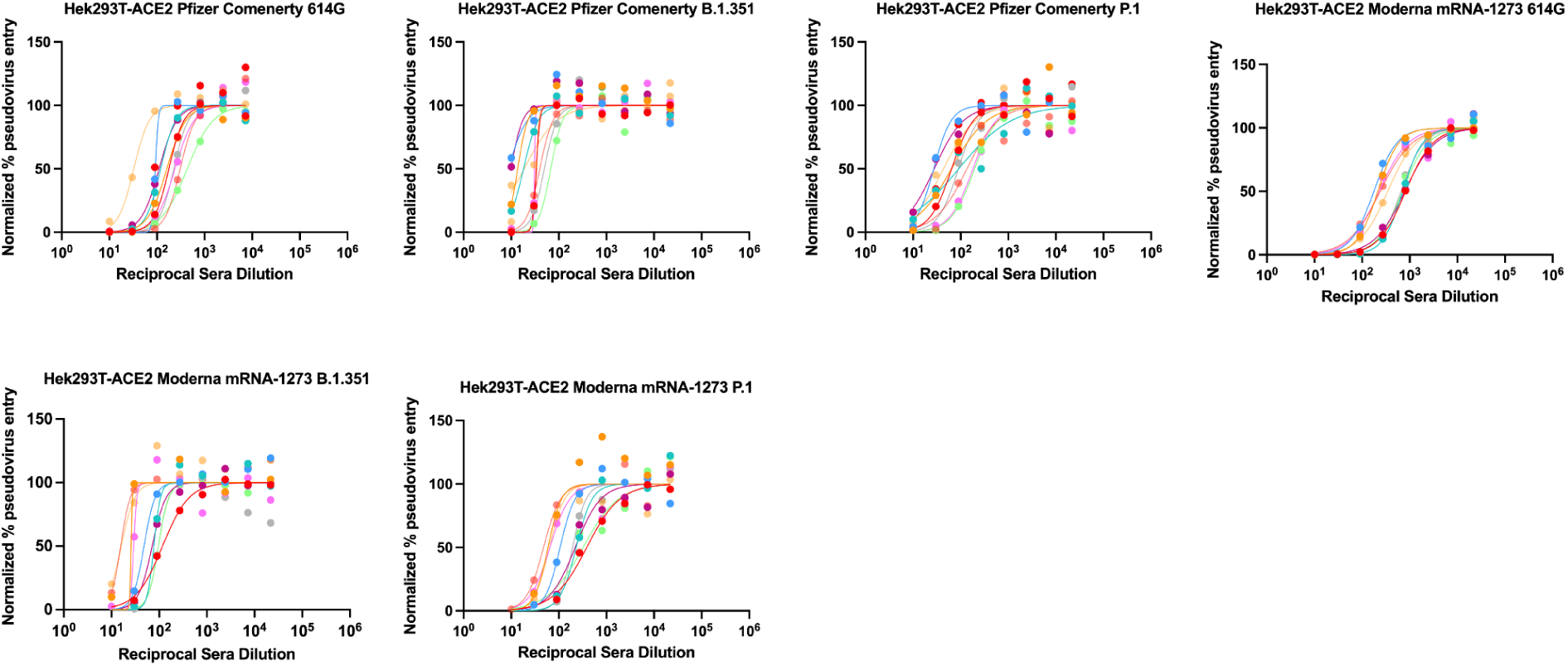
Normalized neutralization curves relating to Figure 1F.

## Methods

### Plasmid construction

The SARS-CoV-2-RBD-Avi construct was synthesized by GenScript into pcDNA3.1-with an N-terminal mu-phosphatase signal peptide and a C-terminal octa-histidine tag, flexible linker, and avi tag (GHHHHHHHHGGSSGLNDIFEAQKIEWHE). The boundaries of the construct are N-_328_RFPN_331_ and _528_KKST_531_-C^24^. The SARS-CoV-2 S ‘2P’ ectodomain trimer (GenBank: YP_009724390.1, BEI NR-52420) was synthesized by GenScript into pCMV with an N-terminal mu-phosphatase signal peptide and a C-terminal TEV cleavage site (GSGRENLYPQG), T4 fibritin foldon (GGGSGYIPEAPRDGQAYVRKDGEWVLLSTPL), and octa-histidine tag (GHHHHHHHH) ^24^. The construct contains the 2P mutations (proline substitutions at residues 986 and 987; and an _682_SGAG_685_ substitution at the furin cleavage site. The SARS-CoV-2 RBD was genetically fused to the N terminus of the trimeric I53-50A nanoparticle component using 16 glycine and serine residues. The RBD-16GS-I53-50A fusion was cloned into pCMV/R using the Xba1 and AvrII restriction sites and Gibson assembly^25^. All RBD-bearing components contained an N-terminal mu-phosphatase signal peptide and a C-terminal octa-histidine tag. SARS-CoV-2 HexaPro construct is as previously described (Hsieh et al.) and placed into CMVR with an octa-his tag.

### Transient transfection

Proteins were produced using endotoxin free DNA in Expi293F cells grown in suspension using Expi293F expression medium (Life Technologies) at 33°C, 70% humidity, 8% CO_2_ rotating at 150 rpm. The cultures were transfected using PEI-MAX (Polyscience) with cells grown to a density of 3.0 million cells per mL and cultivated for 3 days. Supernatants were clarified by centrifugation (5 min at 4000 rcf), addition of PDADMAC solution to a final concentration of 0.0375% (Sigma Aldrich, #409014), and a second spin (5 min at 4000 rcf).

### Microbial protein expression and purification

The I53-50A and I53-50B.4.PT1 proteins^26^ were expressed in Lemo21(DE3) (NEB) in LB (10 g Tryptone, 5 g Yeast Extract, 10 g NaCl) grown in 2 L baffled shake flasks or a 10 L BioFlo 320 Fermenter (Eppendorf). Cells were grown at 37°C to an OD600 ∼ 0.8, and then induced with 1 mM IPTG. Expression temperature was reduced to 18°C and the cells shaken for ∼ 16 h. The cells were harvested and lysed by microfluidization using a Microfluidics M110P at 18,000 psi in 50 mM Tris, 500 mM NaCl, 30 mM imidazole, 1 mM PMSF, 0.75% CHAPS. Lysates were clarified by centrifugation at 24,000 g for 30 min and applied to a 2.6×10 cm Ni Sepharose 6 FF column (Cytiva) for purification by IMAC on an AKTA Avant150 FPLC system (Cytiva). Protein of interest was eluted over a linear gradient of 30 mM to 500 mM imidazole in a background of 50 mM Tris pH 8, 500 mM NaCl, 0.75% CHAPS buffer. Peak fractions were pooled, concentrated in 10K MWCO centrifugal filters (Millipore), sterile filtered (0.22 μm) and applied to either a Superdex 200 Increase 10/300, or HiLoad S200 pg GL SEC column (Cytiva) using 50 mM Tris pH 8, 500 mM NaCl, 0.75% CHAPS buffer. I53-50A elutes at ∼ 0.6 column volume (CV). I53-50B.4PT1 elutes at ∼ 0.45 CV. After sizing, bacterial-derived components were tested to confirm low levels of endotoxin before using for nanoparticle assembly.

### Protein purification

Proteins containing His tags were purified from clarified supernatants via a batch bind method where each clarified supernatant was supplemented with 1 M Tris-HCl pH 8.0 to a final concentration of 45 mM and 5 M NaCl to a final concentration of ∼ 310 mM. Talon cobalt affinity resin (Takara) was added to the treated supernatants and allowed to incubate for 15 min with gentle shaking. Resin was collected using vacuum filtration with a 0.2 μm filter and transferred to a gravity column. The resin was washed with 20 mM Tris pH 8.0, 300 mM NaCl, and the protein was eluted with 3 column volumes of 20 mM Tris pH 8.0, 300 mM NaCl, 300 mM imidazole. The batch bind process was then repeated and the first and second elutions combined. SDS-PAGE was used to assess purity. RBD-I53-50A fusion protein IMAC elutions were concentrated to > 1 mg/mL and subjected to three rounds of dialysis into 50 mM Tris pH 7.4, 185 mM NaCl, 100 mM Arginine, 4.5% glycerol, and 0.75% w/v 3-[(3-cholamidopropyl)dimethylammonio]-1-propanesulfonate (CHAPS) in a hydrated 10K molecular weight cutoff dialysis cassette (Thermo Scientific). S ‘2P’ and HexaPro IMAC elution fractions were concentrated to ∼ 1 mg/mL and dialyzed three times into 50 mM Tris pH 8, 150 mM NaCl, 0.25% L-Histidine in a hydrated 10K molecular weight cutoff dialysis cassette (Thermo Scientific).

### *In vitro* nanoparticle assembly and purification

Total protein concentration of purified individual nanoparticle components was determined by measuring absorbance at 280 nm using a UV/vis spectrophotometer (Agilent Cary 8454) and calculated extinction coefficients. The assembly steps were performed at room temperature with addition in the following order: RBD-I53-50A trimeric fusion protein, followed by additional buffer (50 mM Tris pH 7.4, 185 mM NaCl, 100 mM Arginine, 4.5% glycerol, and 0.75% w/v CHAPS) as needed to achieve desired final concentration, and finally I53-50B.4PT1 pentameric component (in 50 mM Tris pH 8, 500 mM NaCl, 0.75% w/v CHAPS), with a molar ratio of RBD-I53-50A:I53-50B.4PT1 of 1.1:1. All RBD-I53-50 *in vitro* assemblies were incubated at 2-8°C with gentle rocking for at least 30 min before subsequent purification by SEC in order to remove residual unassembled component. Different columns were utilized depending on purpose: Superose 6 Increase 10/300 GL column was used analytically for nanoparticle size estimation, a Superdex 200 Increase 10/300 GL column used for small-scale pilot assemblies, and a HiLoad 26/600 Superdex 200 pg column used for nanoparticle production. Assembled particles were purified in 50 mM Tris pH 7.4, 185 mM NaCl, 100 mM Arginine, 4.5% glycerol, and 0.75% w/v CHAPS, and elute at ∼ 11 mL on the Superose 6 column and in the void volume of Superdex 200 columns. Assembled nanoparticles were sterile filtered (0.22 μm) immediately prior to column application and following pooling of fractions.

### Endotoxin measurements

Endotoxin levels in protein samples were measured using the EndoSafe Nexgen-MCS System (Charles River). Samples were diluted 1:50 or 1:100 in Endotoxin-free LAL reagent water, and applied into wells of an EndoSafe LAL reagent cartridge. Charles River EndoScan-V software was used to analyze endotoxin content, automatically back-calculating for the dilution factor. Endotoxin values were reported as EU/mL which were then converted to EU/mg based on UV/vis measurements. Our threshold for samples suitable for immunization was < 50 EU/mg.

### BALB/c mice for RBD-NP, S ‘2P’, and HexaPro immunizations

Female BALB/c mice (Stock # 000651, BALB/c cByJ mice) four weeks old were obtained from Jackson Laboratory, Bar Harbor, Maine, and maintained at the Comparative Medicine Facility at the University of Washington, Seattle, WA, accredited by the American Association for the Accreditation of Laboratory Animal Care International (AAALAC). Animal procedures were performed under the approvals of the Institutional Animal Care and Use Committee (IACUC) of University of Washington, Seattle, WA.

At six weeks of age, 8 female BALB/c mice per dosing group were vaccinated with a prime immunization, and three weeks later mice were boosted with a second vaccination (IACUC protocol 4470.01). Prior to inoculation, immunogen suspensions were gently mixed 1:1 vol/vol with AddaVax adjuvant (Invivogen, San Diego, CA) to reach a final concentration of 0.01 mg/mL antigen. Mice were injected intramuscularly into the quadriceps muscle of each hind leg using a 27-gauge needle (BD, San Diego, CA) with 50 μL per injection site (100 μL total) of immunogen under isoflurane anesthesia. To obtain sera all mice were bled two weeks after prime and boost immunizations. Blood was collected via submental venous puncture and rested in 1.5 mL plastic Eppendorf tubes at room temperature for 30 min to allow for coagulation. Serum was separated from red blood cells via centrifugation at 2,000 g for 10 min. Complement factors and pathogens in isolated serum were heat-inactivated via incubation at 56°C for 60 min. Serum was stored at 4°C or −80°C until use. The study was repeated twice.

### BALB/c mice for mRNA-1273 immunizations

Female BALB/c mice (6 to 8 weeks old) were obtained from Charles River Laboratories. Animal experiments were carried out in compliance with approval from the Animal Care and Use Committee of Moderna Inc. Mice were immunized with 1µg of mRNA-1273 diluted in 50µl of 1X phosphate-buffered saline (PBS), via intramuscular injection into the same hind leg for both prime and boost. Sera were obtained at 2 (week 5) and 5 (week 8) weeks post-boost for immune analysis. Data are from two independent experiments.All animal procedures done according to the approved SOPs

### K18-hACE2 mice for mRNA-1273 immunizations

K18-hACE2 transgenic mice were purchased from Jackson Laboratories (#034860) and housed in a pathogen-free animal facility at Washington University in St. Louis. Animal studies were carried out in accordance with the recommendations in the Guide for the Care and Use of Laboratory Animals of the National Institutes of Health. The protocols were approved by the Institutional Animal Care and Use Committee at the Washington University School of Medicine (Assurance number A3381-01). mRNA-1273 was produced as described previously^17^. Mice were immunized with 5µg of mRNA-1273 vaccine and boosted with the same dose three weeks later. Serum was obtained three weeks post-boost for immune analysis. Data are from two independent experiments.

### 1292S mice for mRNA-1273 immunizations

129S2 mice (strain: 129S2/SvPasCrl, Cat # 287) were obtained from Charles River Laboratories and housed in a pathogen-free animal facility at Washington University in St. Louis. Animal studies were carried out in accordance with the recommendations in the Guide for the Care and Use of Laboratory Animals of the National Institutes of Health. The protocols were approved by the Institutional Animal Care and Use Committee at the Washington University School of Medicine (Assurance number A3381-01). Mice were immunized with 5µg of mRNA-1273 vaccine and boosted with the same dose three weeks later. Serum was obtained three weeks post-boost for immune analysis. Data are from two independent experiments.

### Pigtail macaques

Two adult male Pigtail macaques (*Macaca nemestrina*) were immunized in this study. All animals were housed at the Washington National Primate Research Center (WaNPRC), an AAALAC International accredited institution. All experiments were approved by The University of Washington’s IACUC. Animals were singly housed in comfortable, clean, adequately-sized cages with ambient temperatures between 72–82°F. Animals received environmental enrichment for the duration of the study including grooming contact, perches, toys, foraging experiences and access to additional environment enrichment devices. Water was available through automatic watering devices and animals were fed a commercial monkey chow, supplemented daily with fruits and vegetables. Throughout the study, animals were checked twice daily by husbandry staff.

Two Pigtail macaques were immunized with 250 μg of RBD-12GS-I53-50 nanoparticle (88 μg RBD antigen) with AddaVax at day 0 and day 28. Blood was collected every 14 days post-prime. Blood was collected in serum collection tubes and allowed to clot at room temperature. Serum was isolated after a 15 min spin at 1455 x g for 15 min and stored at −80°C until use. Prior to vaccination or blood collection, animals were sedated with an intramuscular injection (10 mg/kg) of ketamine (Ketaset®; Henry Schein). Prior to inoculation, immunogen suspensions were gently mixed 1:1 vol/vol with AddaVax adjuvant for immunizations 1 and 2 and O/W for immunization 3 (Invivogen, San Diego, CA) to reach a final concentration of 0.250 mg/mL antigen. The vaccine was delivered intramuscularly into both quadriceps muscles with 1 mL per injection site on days 0 and 28. All injection sites were shaved prior to injection. Animals were observed daily for general health (activity and appetite, urine/feces output) and for evidence of reactogenicity at the vaccine inoculation site (swelling, erythema, and pruritus) for up to 1 week following vaccination. They also received physical exams including temperature and weight measurements at each study time point. None of the animals became severely ill during the course of the study nor required euthanasia.

### Rhesus macaques

Rhesus macaques (Macaca mulatta) of Indian origin, aged 3–7 years were assigned to the study^1^. Animals were distributed between the groups such that the age and weight distribution were comparable across the groups. Animals were housed and maintained at the New Iberia Research Center (NIRC) of the University of Louisiana at Lafayette, an AAALAC International accredited institution, in accordance with the rules and regulations of the Guide for the Care and Use of Laboratory Animal Resources. The entire study (protocol 2020-8808-15) was reviewed and approved by the University of Louisiana at Lafayette IACUC. All animals were negative for SIV, simian T cell leukemia virus, and simian retrovirus.

Adapted from ref. ^1^: AS03 was kindly provided by GSK Vaccines. AS03 is an oil-in-water emulsion that contains 11.86 mg a-tocopherol, 10.69 mg squalene, and 4.86 mg polysorbate 80 (Tween-80) in PBS. For each dose, RBD-NP was diluted to 50 μg/ml (RBD component) in 250 μl of Tris-buffered saline (TBS) and mixed with an equal volume of AS03.The dose of AS03 was 50% v/v (equivalent of one human dose). Soluble HexaPro was diluted to 50 μg/ml in 250 μl of Tris-buffered saline (TBS) and mixed with an equal volume of AS03. All immunizations were administered via the intramuscular route in right forelimbs. The volume of each dose was 0.5 ml.

### Pfizer and Moderna vaccinated human sera

Sera samples were collected from participants who had received both doses of the Pfizer or Moderna mRNA vaccine and were 7–30 days post second vaccine dose. Clinical data for these individuals are summarized in **Table S1**. Individuals were enrolled in the UWARN: COVID-19 in WA study at the University of Washington in Seattle, WA. This study was approved by the University of Washington Human Subjects Division Institutional Review Board (STUDY00010350).

### Pseudovirus production

G614 SARS-CoV-2 S (YP 009724390.1), B.1.351 S, and P.1 S pseudotyped VSV viruses were prepared as described previously^11,27,28^. Briefly, HEK293T cells in DMEM supplemented with 10% FBS, 1% PenStrep seeded in 10-cm dishes were transfected with the plasmid encoding for the corresponding S glycoprotein using lipofectamine 2000 (Life Technologies) following manufacturer’s indications. One day post-transfection, cells were infected with VSV(G*ΔG-luciferase)^29^ and after 2 h were washed five times with DMEM before adding medium supplemented with anti-VSV-G antibody (I1-mouse hybridoma supernatant, CRL-2700, ATCC). Virus pseudotypes were harvested 18-24 h post-inoculation, clarified by centrifugation at 2,500 x g for 5 min, filtered through a 0.45 μm cut off membrane, concentrated 10 times with a 30 kDa cut off membrane, aliquoted and stored at −80°C.

### Pseudovirus Neutralization

HEK293-hACE2 cells^15^ or VeroE6-TMPRSS2^14^ were cultured in DMEM with 10% FBS (Hyclone) and 1% PenStrep and 8ug/mL puromycin for TMPRSS2 maintenance with 5% CO_2_ in a 37°C incubator (ThermoFisher). One day prior to infection, 40 μL of poly-lysine (Sigma) was placed into 96-well plates and incubated with rotation for 5 min. Poly-lysine was removed, plates were dried for 5 min then washed 1 × with water prior to plating with 40,000 cells. The following day, cells were checked to be at 80% confluence. In an empty half-area 96-well plate a 1:3 serial dilution of sera was made in DMEM and diluted pseudovirus was then added to the serial dilution and incubated at room temperature for 30-60 min. After incubation, the sera-virus mixture was added to the cells at 37°C and 2 hours post-infection, 40 μL 20% FBS-2% PenStrep DMEM was added. After 17-20 hours VSV 40 μL/well of One-Glo-EX substrate (Promega) was added to the cells and incubated in the dark for 5-10 min prior reading on a BioTek plate reader. Measurements were done in at least duplicate. Relative luciferase units were plotted and normalized in Prism (GraphPad). Nonlinear regression of log(inhibitor) versus normalized response was used to determine IC_50_ values from curve fits. Kruskal Wallis tests were used to compare two groups to determine whether they were statistically different. Fold changes were determined by comparing individual animal IC_50_ and then averaging the individual fold changes for reporting.

### Pseudovirus Neutralization Assay for BALB/c mRNA-1273 samples

Codon-optimized full-length spike genes (Wuhan-Hu-1 with G614, Beta, or Gamma) were cloned into pCAGGS vector. Spike genes of Beta and Gamma variants contain the following mutations: L18F-D80A-D215G-L242-244del-R246I-K417N-E484K-N501Y-D614G-A701V, and L18F-T20N-P26S-D138Y-R190S-K417T-E484K-N501Y-D614G-H655Y-T1027I-V1176F, respectively. To generate VSVΔG-based SARS-CoV-2 pseudovirus, BHK-21/WI-2 cells were transfected with the spike expression plasmid and infected by VSVΔG-firefly-luciferase as previously described^30^. A549-hACE2-TMPRSS2 cells were used as target cells for the neutralization assay. Lentivirus encoding hACE2-P2A-TMPRSS2 was made to generate A549-hACE2-TMPRSS2 cells which were maintained in DMEM supplemented with 10% fetal bovine serum and 1µg/mL puromycin. To perform neutralization assay, mouse serum samples were heat-inactivated for 45 minutes at 56 Celsius and serial dilutions of the samples were made in DMEM supplemented with 10% fetal bovine serum. The diluted serum samples or culture medium (serving as virus only control) were mixed with VSVΔG-based SARS-CoV-2 pseudovirus and incubated at 37 Celsius for 45 minutes. The inoculum virus or virus-serum mix was subsequently used to infect A549-hACE2-TMPRSS2 cells for 18 hr at 37 Celsius. At 18 hr post infection, equal volume of One-Glo reagent (Promega; E6120) was added to culture medium for readout using BMG PHERastar-FS plate reader. The percentage of neutralization is calculated based on relative light units of the virus control, and subsequently analyzed using 4 parameter logistic curve (Prism 8).

### Live virus focus-reduction neutralization test for K18-hACE2 and 1292S mRNA-1273 samples

SARS-CoV-2 G614 was produced by introducing the mutation into an infectious clone of WA1/2020^31^. B.1.351 was isolated from an infected individual. Both viruses were propagated on Vero-TMPRSS2 cells and subjected to deep sequencing to confirm the presence of expected substitutions. Focus-reduction neutralization tests (FRNTs) were performed as described^32^. Briefly, serial dilutions of antibody were incubated with 10^2^ FFU of SARS-CoV-2 for 1 h at 37°C. Immune complexes were added to VeroE6-TMPRSS2 cell monolayers and incubated for 1 h at 37°C prior to the addition of 1% (w/v) methylcellulose in MEM. Following incubation for 30 h at 37°C, cells were fixed with 4% paraformaldehyde (PFA), permeabilized and stained for infection foci with an oligoclonal mixture of ati-SARS-CoV-2 (SARS2-02, SARS2-11, SARS2-31, SARS2-38, SARS2-57, and SARS2-71^33^; diluted to 1 mg/mL total mAb concentration). Antibody-dose response curves were analyzed using non-linear regression analysis (with a variable slope) (GraphPad Software). The antibody half-maximal inhibitory concentration (ID50) required to reduce infection was determined.

### CoV database parameters

Data sheets were downloaded on December 1st, 2021 from https://covdb.stanford.edu/page/susceptibility-data/ selecting for either Pfizer BNT162b2 or Moderna mRNA1273 plasma Abs and either B.1.351 or P.1 variant and using the median fold change reported metric. Data sets where individuals were previously infected or had only 1 shot were excluded. Data sets where the fold-change comparator was not a Wuhan-Hu-1 derivative (WA1, G614) or datasets where the number of samples measured was below 3 were excluded.

